# Paradigm Shift in Sensorimotor Control Research and Brain Machine Interface Control: The Influence of Context on Sensorimotor representations

**DOI:** 10.1101/239814

**Authors:** Yao Zhao, John P. Hessburg, Jaganth Nivas Asok Kumar, Joseph Thachil Francis

**Affiliations:** Joint Program in Biomedical Engineering, Polytechnic Institute of NYU and SUNY Downstate, Brooklyn, NY; Department of Physiology and Pharmacology, State University of New York Downstate Medical School, Robert F Furchgott Center for Neural and Behavioral Science, Brooklyn, NY; Department of Biomedical Engineering, Cullen College of Engineering, University of Houston, Houston, TX

**Keywords:** Reinforcement learning, Brain Machine Interface (BMI), Motor Cortex, Somatosensory Cortex, Dopamine, Sensorimotor control

## Abstract

Neural activity in the primary motor cortex (M1) is known to correlate with movement related variables including kinematics and dynamics. Our recent work, which we believe will lead to a paradigm shift in sensorimotor research, has shown that in addition to these movement related variables, activity in M1, and the primary somatosensory cortex (S1), are also modulated by context, such as value, during both active movement and movement observation. Here we expand on the investigation of reward modulation in M1, showing that reward level changes the neural tuning function of M1 units to both kinematic as well as dynamic related variables. In addition, we show that this reward-modulated activity is present during brain machine interface (BMI) control. We suggest that by taking into account these context dependencies of M1 modulation, we can produce more robust BMIs. Toward this goal, we demonstrate that we can classify reward expectation from M1 on a movement-by-movement basis under BMI control and use this to gate multiple linear BMI decoders toward improved offline performance. These findings demonstrate that it is possible and meaningful to design a more accurate BMI decoder that takes reward and context into consideration. Our next step in this development will be to incorporate this gating system, or a continuous variant of it, into online BMI performance

## 1 Introduction

Primary motor cortical (M1) activity appears to encode movement related kinematics and dynamics, and is often modeled as a linear relationship between neuronal firing and desired movement toward the development of brain-machine interfaces (BMIs) (Chhatbar & Francis 2013). BMIs allow subjects to control physical or virtual systems including robotic arms and computer cursors using neural signals (Li et al 2011, Serruya et al 2002, Taylor et al 2002, Velliste et al 2008). BMIs have been used to restore reaching and grasping for paralyzed patients with some success (Ajiboye et al 2017, Bouton et al 2016, Collinger et al 2013, Hochberg et al 2012), and making such systems more robust and easier to control is of great importance. A critical component of BMIs are the motor cortex tuning models, which describe the relationship between neural firing rates and kinematics, such as hand/endpoint velocity (Taylor et al 2002), or dynamics, such as force and torques (Carmena et al 2003, Chhatbar & Francis 2010, Chhatbar & Francis 2013, Suminski et al 2011).

Recently our lab and others have shown that context can modulate the primary motor cortex (M1) (Downey et al 2017, Marsh et al 2015, Ramakrishnan et al 2017, Ramkumar et al 2016). For instance, it has become clear that a reward signal exists in the primary motor cortex (M1), and primary somatosensory cortex (S1) as well (McNiel et al 2016a, McNiel et al 2016b). We hypothesized (Marsh et al 2015) that this reward signal originates in midbrain dopaminergic areas such as the ventral tegmentum (VTA) and substantia nigra pars compacta (SNc), as there are dopaminergic receptors in M1 as well as terminals from these dopaminergic centers (Richfield et al 1989). Dopamine is necessary for LTP in M1 (Molina-Luna et al 2009), and has been shown to have a “charging” effect on neural activity, possibly acting as a motivational signal (Hamid et al 2016, Hollerman & Schultz 1998, Schultz 2000). Although the aforementioned studies showed that M1 neurons multiplex the representation of reward and motor activity, it has yet to be concretely characterized whether such reward modulation could be used to design a more robust and accurate BMI decoder. In the current study, we introduce the fact that reward modulates neural activity related to dynamic variables, such as grip force, and to BMI controlled kinematic variables for the first time to our knowledge.

The current work has two main goals. First, to show that significant differences exist in both directional and force tuning models of M1 units between rewarding and non-rewarding trials, and second, that reward level (Tarigoppula et al 2017), that is the value of a given movement, can be used as additional information to improve BMI decoding accuracy, at least in the offline, open-loop system.

## 2 Methods

### 2.1 Surgery

Our subjects were non-human primates (NHPs), one male rhesus macaque (monkey S) and one female bonnet macaque (monkey P) were implanted in M1 with chronic 96-channel platinum microelectrode arrays (Utah array, 10*10 array separated by 400 μm, 1.5 mm electrode length, ICS-96 connectors, Blackrock Microsystems). The hand and arm region of M1 contralateral to their dominant hand was implanted with the same technique as our previous work (Chhatbar et al 2010, Marsh et al 2015). All surgical procedures were conducted in compliance with guidelines set forth by the National Institutes of Health Guide for the Care and Use of Laboratory Animals and were approved by the State University of New York Downstate Institutional Animal Care and Use Committee. Briefly, general anesthesia was administered, and aseptic conditions were maintained throughout the surgery. Animal preparation and anesthesia were performed directly or supervised by members of the State University of New York Downstate Division of Comparative Medicine veterinary staff. Ketamine was used to induce anesthesia, and isofluorane and fentanyl were used for maintenance. Dexamethasone was used to prevent inflammation during the procedure and diuretics such as mannitol and furosemide, were available to further reduce cerebral swelling if needed. Both subjects were observed hourly for the first 12 hours after implantation and were provided with a course of antibiotics (baytril and bicilin) and analgesics (buprenorphine and rimadyl).

### 2.2 Extracellular unit recordings

After a 2-3 week recovery period, spiking and local field potential (LFP) activities were simultaneously recorded with multichannel acquisition processor systems (MAP, Plexon Inc.) while the subjects performed the experimental task. Neural signals were amplified, bandpass filtered (170 Hz to 8 kHz for single and multiunit activity and 0.7–300 Hz for LFPs), sampled at 40 kHz for single-unit/ multi-unit activity, 2 KHz for LFP, and manually thresholded for single units. Single and multi-units were sorted based on their waveforms using principal component (PC)-based methods in Sort-Client software (Plexon Inc).

### 2.3 Behavioral task

Monkeys S and P were trained to perform a reach-grasp-transport-release task depicted in Figure 1. In this task, the subjects controlled certain aspects of a simulated anthropomorphic robotic arm (Barrett WAM) in order to reach-grasp-transport-release target cylinders. Each trial consisted of 6 stages: cue display, reaching, grasping, transporting, releasing and reward delivery. At the start of a trial, cues were displayed (green squares) to indicate the level of reward, juice, the animal would receive upon successful completion of the task. The number of green squares (0 - 3) corresponded to the number of 0.5 second juice delivery periods. If no green square was displayed, then no reward was delivered upon successful completion of the trial. In an unsuccessful trial, no reward was delivered and the trial was repeated at the same reward level until completed successfully, thus motivating the subjects to complete even the zero reward level trials successfully. Two NHPs conducted two sessions of a manual grip force control version of the above task as well as two sessions each of a BMI version of the task.

**Figure 1.**
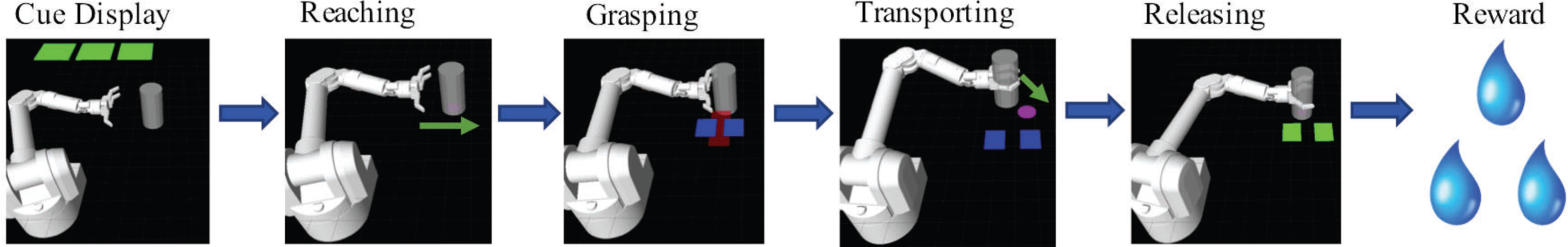
Behavioral Task: The behavioral task was composed of 6 scenes. First, during the cue display scene the animal was cued via the number of green squares to the amount of reward it would receive if it completed a trial successfully. Each green square indicated 0.5 seconds worth of liquid reward. Zero green squares indicated a non-rewarding trial. Trials could either be under manual control or BMI control. In manual control the NHP had to output the cued, blue rectangles, grip force, red rectangle, in order for the trial to be successful. In BMI control mode, the NHP had to control the reaching trajectory of the task (see methods).

For the manual task, the virtual arm reached the target cylinder automatically. Then the animal controlled the grasping motion of the hand by manually squeezing a force transducer with its dominant hand. The amount of force applied was represented in the virtual environment by a red rectangle that increased in width proportional to the force output. The subject had to reach and maintain a level of force indicated by a pair of blue force target rectangles (fig.1). The robotic arm then automatically moved the cylinder to a target location while the animal maintained the target grip force. When the arm reached the target location the animal released the gripper, placing the cylinder at the target location, resulting in a successful trial. On trials cued as rewarding via the green squares, the animal received a juice reward on successful completion of the trial. If non-rewarding cue, zero green squares, was presented, or if the animal was unsuccessful in completing the trial, no reward was delivered and the task moved to the next trial, which was of the same value as the previous unsuccessful trial. After reward delivery, the arm retreated horizontally left automatically to the starting position.

For the BMI task, after the reward cue presentation, the subject controlled the robotic arm’s movement via the BMI in order to reach towards the target cylinder using M1 activity. During the reaching stage of the task, the cylinder was always located horizontally to the right from the starting position of the virtual hand. When within a threshold distance, the cylinder was grasped automatically. The animal then needed to move the cylinder to the target position using BMI control. The location of potential targets was random in the 2D plane. If the animal brought the cylinder to the target position, the trial was successful. The hand automatically released the grasp on the cylinder, and then the arm reset to the starting position automatically.

This study was designed to investigate the effect of varying levels of reward on neural encoding, there were two versions for both BMI and the manual task differing in the reward levels offered during a session. The reward level was determined by the amount of time the juice reward straw (electronically controlled by a solenoid) was kept open. The solenoid was opened for 0.5 seconds for every successive level of reward. For the first recording session, the two reward levels were 0 (non-rewarding) and 1 (0.5 seconds of reward delivery). For the second session, the reward levels were 0 and 3 (1.5 seconds of reward delivery). Further analysis has been conducted on both single and multi-unit data. For the manual task, 77 M1 units were recorded from Monkey S and 64 M1 units were recorded from Monkey P. For the BMI task, 87 M1 units were recorded from Monkey S and 102 M1 units were recorded from Monkey P.

During BMI trials, subjects controlled the virtual arm movement during the reaching and transporting stages. The BMI decoder used was a ReFIT Kalman filter (Gilja et al 2012), which used binned firing rates (100 ms bins) to predict the animal’s command for the virtual arm’s velocities. The Kalman filter is a linear dynamic system, which has a state model and observation model.

At time *t*, the state model is:

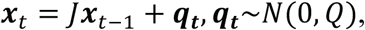

and the observation model is:

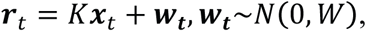

where **bold**, will be used to represent vectors, such as ***x***_t_, which is the state vector representing positions and velocities at time bin *t*, ***r***_t_ is the observation variable representing binned firing rates at time bin *t*, and ***q***_**t**_ and ***w***_t_ are Gaussian noise. Given *J*, *Q*, *K*, and *W*, the Kalman filter can provide the best estimation for the current state ***x***_t_ based on the previous state ***x***_**t−1**_ and the current observation ***r***_t_. The ReFIT Kalman filter allows one to retrain and improve parameters *J*, *Q*, *K*, *W*, using intended velocities during previous BMI control (Gilja et al 2012). The bin size used here was 100 ms. The Re-fit Kalman decoder has been retrained every block (8000 time bins, 800 second).

An assistive controller (Fig. 2) was used to manage frustration levels of the NHP subjects on the BMI task and allowed us to change the difficulty of the task at will. In assistive control, the velocity commands *v*_*cxt*_ and *v*_*cyt*_ that controlled the virtual arm’s movement in the x and y dimensions were a linear combination (H) of BMI decoded velocities *v*_*x1t*_ and *v*_*y1t*_ and “intended” velocities that we will simply refer to as *v*_*xt*_ and *v*_*yt*_ as they are used for most of what comes below. That linear combination (H) is:

**Figure 2.**
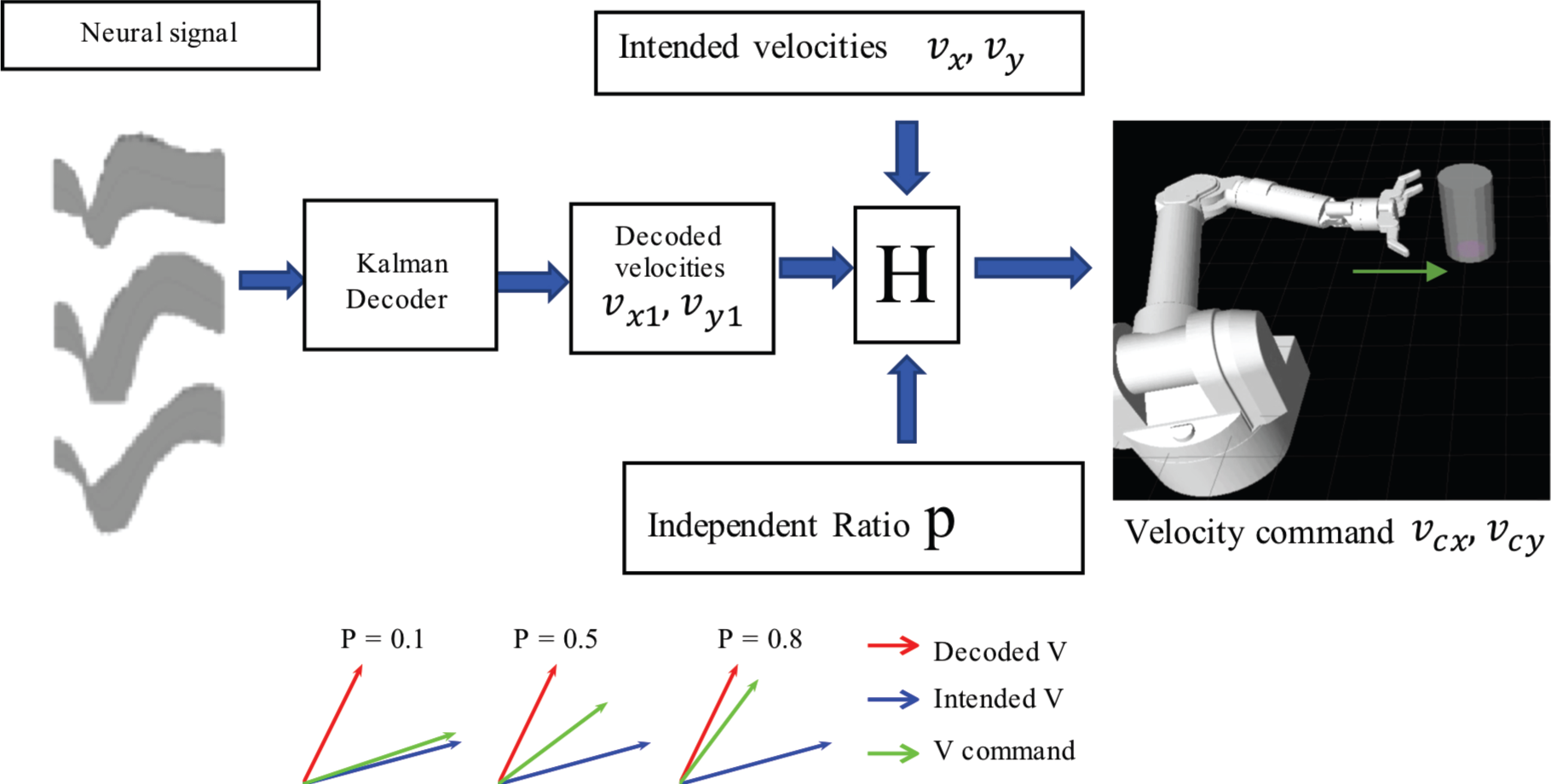
Assistant BMI control. Velocity command is the linear combination of predicted velocities (output from the ReFIT Kalman decoder) and intended velocities. The velocity of the virtual hand is a linear combination (H) of ReFIT Kalman decoder output and the intended velocity. Higher the p is, the velocity command is closer to the decoder output.

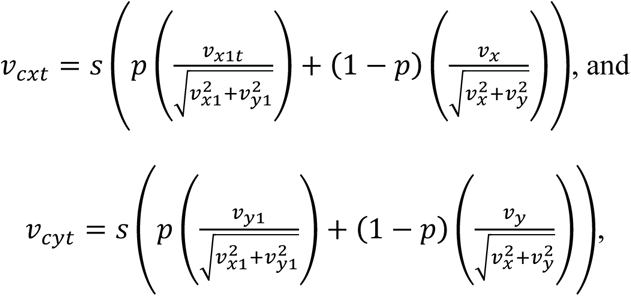

where *s* is a constant speed. We manually set *s* = 40*cm*/*s*. Animals reached the cylinder in about 0.5s at that speed.

We define intended velocities as in the direction of the target location, but with the speed of the decoded velocities, see below. Thus, the difficulty of the task could be changed by adding more or less of the intended velocity by adjusting the independence ratio P. A higher value of P indicates that the velocity command relies more on the BMI decoder output than the intended velocities. All BMI data used here was recorded in sessions with an independence ratio greater than 0.8.

### 2.4 Off-line data analysis

#### 2.4.1 Tuning curve analysis

Linear regression was performed to fit neural encoding models for both the manual task and the BMI task. For the BMI task, during the reaching and transporting scenes the linear intended velocity-encoding model was given by:

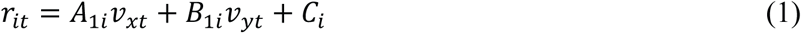

where *r*_*it*_ is unit i’s binned firing rate at time bin *t* (100 ms bins). Again, *v*_*xt*_ and *v*_*yt*_ are the BMI controlled virtual hand’s intended velocities at time bin *t* in the x and y directions, respectively. The intended velocity at time bin *t* has the same speed as the real BMI controlled velocity at time bin *t*, but the direction is towards the target (Gilja et al 2012). If considering movement direction rather than velocity, we have:

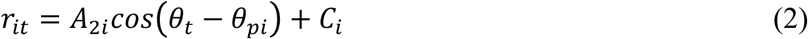

where *θ*_*t*_ indicates the intended movement direction, which is the direction towards the target at time bin *t*. *θ*_*pi*_ is the preferred direction of the i^th^ unit, defined as the direction of movement that evoked the maximum firing rate in that unit. Equation (2) is referred to as the intended directional tuning curve equation and can be used for BMI decoding (Moran and Schwartz, 1999). Equation (2) and equation (1) are equivalent. Let:

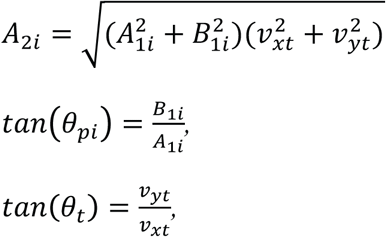

Then equation (1) can be related to equation (2).

Intended velocities and directions were used instead of real velocities and directions for fitting equations (1) and (2) because it has been previously shown that ReFIT Kalman decoder using intended kinematics information can correct model parameters and the decoder has better BMI performance (Gilja et al 2012).

Considering rewarding and non-rewarding trials separately, the following equations can be written:

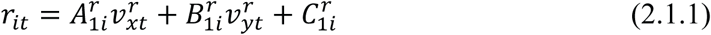

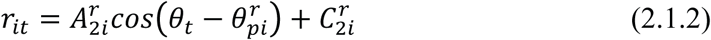

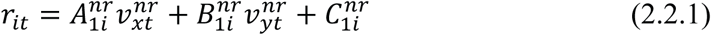

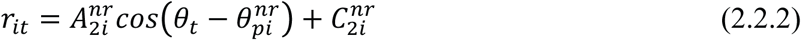

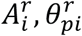 and 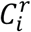 are parameters for rewarding trials (r), and 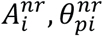 and 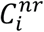 are parameters for non-rewarding trials (nr). Equation (2.1.1) and (2.2.1) were fit for all units using Matlab function “regstats”. Units were considered significant if (2.1.1) and (2.2.1)differed (p<0.05) from a constant model, that is 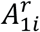 or 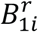 from equation (2.1.1) and 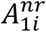 or 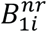 were significant different from zero (p<0.05). Equation (2.1.2) was fit for all units using data from all rewarding trials during the transport stage (free 2D movement). Similarly, equation (2.2.2) was fit for all units using data from all non-rewarding trials during the transport stage. Both equations were fit using a nonlinear least square fit. To minimize overfitting, data from the reaching stage was not included for analysis as all the reaching movements were the same, being made horizontally towards the right from the starting position. All estimated values and variances for parameters 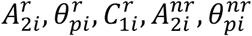 and 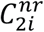 from (2.1.2) and (2.2.2) were calculated using non-linear least squares fit in Matlab. If any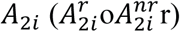 was less than zero, then the value was multiplied by −1 to yield a positive value of *A*_2i_, and the corresponding *θ*_*pi*_was adjusted by:

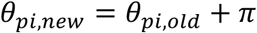

After this adjustment, *θ*_*pi*_ would always represent the preferred direction of the unit, which means that unit have maximum firing rate when the moving direction is *θ*_*pi*_.

For each unit, the shapes of the two tuning curves defined by equations (2.1) and (2.2) could then be compared. T-tests were conducted to compare 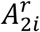 and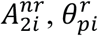 and 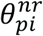 for every unit to see if the amplitude or preferred direction were significantly different. For every unit that had a significant difference between either amplitudes or preferred directions between rewarding and non-rewarding trials, the difference between two preferred directions ∆*θ*_*pi*_ and the normalized difference between two amplitudes ∆*A*_*i*_ were calculated:

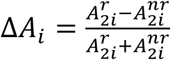

∆*θ*_*pi*_ was the angle between two unit vectors 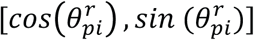, and 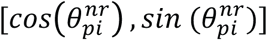.

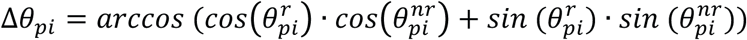

By this definition, ∆*θ*_*pi*_ was in the range of [0, π].

For the manual task, during the grasping, transporting, and releasing scenes, the linear force encoding model was given by:

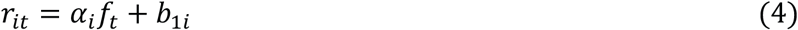

where *r*_*it*_ is the *ith* unit’s binned firing rate at time bin *t*, and *f*_*t*_ is the grip force at time bin *t*. Similarly, to the previous analysis, equation (4) for all units was fit with the Matlab function “regstats” to find all significant units (p<0.05). For every significant unit, the linear force encoding models which consider rewarding and non-rewarding trials separately are:

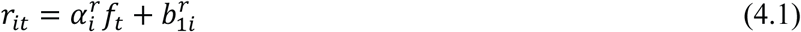

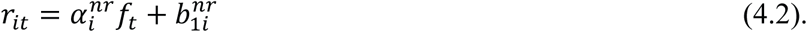

*α*_*ir*_ (rewarding) and *α*_*inr*_ (non-rewarding) for each unit were then compared using one-way analysis of covariance (ANCOVA) to see if the two slopes had a significant difference (p<0.05). For each unit where the two slopes were significantly different, the normalized difference between the slopes ∆*α*_*i*_ was calculated:

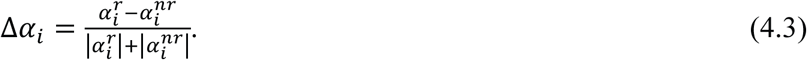

When plotting example tuning curves, normalized firing rates *r*_*norm*_ were calculated:

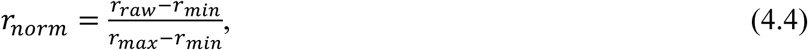

where *r*_*raw*_ is the raw firing rate, *r*_*max*_ is the maximum firing rate though all time in the same block (for both rewarding and non-rewarding trials) and *r*_*min*_ is the minimum firing rate for that unit.

#### 2.4.2 Linear decoding model considering reward level

From these neural encoding models of rewarding and non-rewarding trials, a combined linear decoding/prediction kinematics model was designed that took multiple reward levels into consideration (decoder 2). For the velocity decoder, the decoding accuracies between decoder 1, where all trials were considered together, and decoder 2, treating rewarding and non-rewarding trials separately, were compared using 5-fold cross validation. For velocity decoder 1, the linear decoding model is:

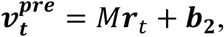

For velocity decoder 2, the linear decoding models are:

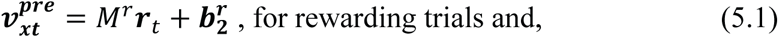

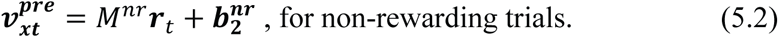

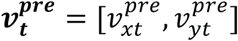′ is the decoded/predicted velocity at time bin *t*, ***r***_*t*_ is the *n*-dimensional firing rate vector at time bin *t*, where *n* is the total number of units. For each testing data set, the velocity error at time bin *t* is:

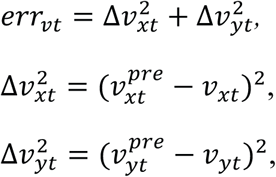

where 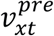 and 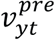 are predicted velocities at time bin t and *v*_*xt*_ and *v*_*yt*_ are intended velocities at time bin *t*. For the single linear decoder (decoder 1), the total error is given by:

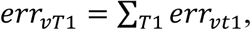

where *T*_1_ is total time bins for velocity decoder 1, which is the total number of time bins for all trials, and *err*_*vt1*_ is the velocity error at time bin *t* using decoder 1. For velocity decoders 2.1 and 2.2, we have:

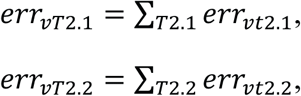

where *T*_2.1_ is total time bins for velocity decoder 2.1 (all rewarding trials), *T*_2.2_ is total time bins for velocity decoder 2.2 (all non-rewarding trials), *err*_*vt*2.1_ is the velocity error at time bin *t* using decoder 2.1, and *err*_*vt*2.1_ is the velocity error at time bin *t* using decoder 2.2. Since *T*_1_ = *T*_2.1_ + *T*_2.2_, *p*_*ve*_ is defined to quantify the percent error reduction:

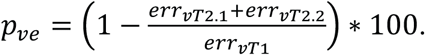

*p*_*ve*_ can be used to compare the velocity decoding accuracy of decoder 1 and decoder 2.

Similarly, for the force decoder the decoding accuracies between decoder 1 and 2 were also compared using 5-fold cross validation.

For force decoder 1, the linear decoding model is:

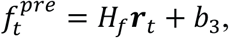

For force decoder 2, the linear decoding models are:

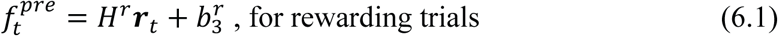

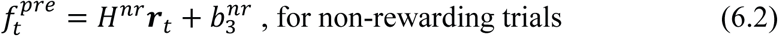

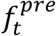 is the decoded force at time bin *t*, ***r***_t_ is the firing rate vector at time bin *t*, ***r***_t_ is an *n*-dimensional vector, and *n* is the total number of units. For each testing data set, the force error at time bin *t* is:

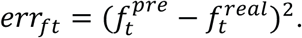

where 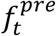 is the decoded/predicted force at time bin *t* and 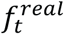 is real force at time bin *t*. The total error for force decoders 1, 2.1 and 2.2 are given by:

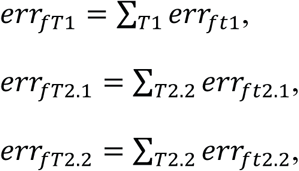

where *T*_1_ represents all time bins for force decoder 1 over all trials, *T*_2.1_ represents all time bins for force decoder 2.1 over all rewarding trials, *T*_2.2_ represents all time bins for force decoder 2.2 over all non-rewarding trials, *err*_*ft1*_ is the force error at time bin *t* using decoder 1, *err*_*ft2.1*_ is the force error at time bin *t* using decoder 2.1, and *err*_*ft2.2*_ is the force error at time bin *t* using decoder 2.2. Since *T*_1_ = *T*_2.1_ + *T*_2.2_, *p*_*fe*_ is defined as:

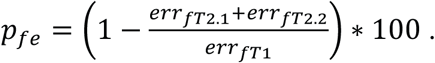

To make sure the error reduction did not because of decoder 2 had more parameters than decoder 1, control groups have been built. We shuffled the reward label for each trail randomly then rand decoder 2 again. This way we still have the same two separately linear decoders, but the separation was random. *p*_*ve*_ or *p*_*fe*_ for that random shuffled decoder 2 were calculated using the same method above. Marked as *p*_*ves*_ or *p*_*fes*_. That shuffles have been done for 500 times in each data block, which means for each data block, we have 500 *p*_*ves*_ or *p*_*fes*_ samples. For each block, we compared its *p*_*ve*_ (for BMI task) or *p*_*fe*_ (for manual task) and its corresponding *p*_*ves*_ or *p*_*fes*_ distribution. We first tested if *p*_*ves*_ or *p*_*fes*_ distribution was Gaussian using Jarque-Bera test (p<0.05) using Matlab function “JBtest”, then compared if the *p*_*ve*_ or *p*_*fe*_ from a certain block was greater than its corresponding 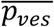 or 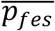, and if that difference was significant using t-test. That control groups have been built for both BMI and manual task.

#### 2.4.3 Classification algorithm

Previous results show that post-cue firing rates in M1 are separable between rewarding and non-rewarding trials (Marsh et al 2015). A k-nearest neighbors (kNN) algorthm was used as a classifier, with firing rates from 0.3 seconds to 0.9 seconds (6 time bins) after the cue as the input. The kNN algorithm was chosen because it is a nonparametric algorithm that allows for a nonlinear decision boundary and also has an acceptable computational complexity for a small sample size such as the one used in this study, which has less than 200 samples (Altman 1992). The time period from 0.3s to 0.9s was chosen because for the manual task, this is the time period when the virtual arm is horizontally moving right to reach the cylinder. For the BMI task, this is the time period when the virtual arm is moving horizontally right from the start position to to the cylinder position. Since the virtual arm movement direction is always the same for that time period in the manual task and the cylinder position was always the same from the start position for the BMI task, the neural movement encoding is simlar among trials. Thus, this time period may show a clear separability between rewarding and non-rewarding trials.

For trial *t*, the input is the firing rate vector ***R***_t_. ***R***_t_ is the 6*n*-dimensional firing rate vector at trial *t*. Each element in ***R***_t_ represents one unit’s firing rate for each of the six bins in the above mentioned time period. The output is the class label ***l***_t_ where ***l***_t_ = 0 for non-rewarding and ***l***_t_ = 1 for rewarding trials. For any testing data ***R***_*test*_, the 5 nearest neighbors (Euclidean distance) of ***R***_*test*_ were found in training data using euclidean distance and named as ***R***_*n*1_~***R***_*n*5_. We then obtain:

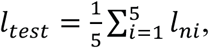

where *l*_ni_ are the labels for ***R***_**ni**_, *i* = 1,2,3,4,5 respectively. The test trial was classified as rewarding if ***l***_test_ > 0.5, and classified as non-rewarding if ***l***_test_ < 0.5.

Not all M1 units have reward modulation, so we hypothesized that a better classifier could be built by using the subset of units which show reward modulation. To choose this optimal subset, the best individual unit ensemble construction procedure (Leavitt et al 2017a) was used. This consists of: 1) building classifiers using each individual unit, 2) ranking all units based on the classification accuracy of that unit’s classifier, and 3) iteratively adding ordered units to make a more complex classifier at each iteration. If the new unit added reduced the overall classification accuracy, it was dropped from consideration. The final classifier therefore used only the units that were beneficial to it, and did not use the remainder of the units.

#### 2.4.4 Two-stage decoder

Combining the post-cue classifier (method 1.4.3) and the separate linear model (method 1.4.2), a two-stage decoder was designed to incorporate multiple reward levels. The first stage is the kNN classifier (method 1.4.3), using post-cue firing rates at the beginning of each trial to determine the reward level for the trial. The second stage then consists of the two different linear decoders obtained from stage one. The second-stage decoder’s output is velocity (using equations (5.1) and (5.2)) for the BMI task or force (using equations (6.1) and (6.2)) for the manual task. Using this two-stage decoder, the reward level can be calculated directly from the population firing rates, and no additional information is needed for the linear decoders in the second stage.

An offline test was run for the two-stage decoder, and its decoding accuracy was compared to a single stage linear decoder. *err*_*vt*_and *err*_*ft*_ were calculated using the same equations as described previously. *p*_*ve*_ and *p*_*fe*_ for the two-stage decoder were defined as:

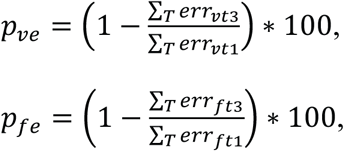

where *T* represents the total time bins, *err*_*vt3*_ is the velocity prediction error at time bin *t* using the two-stage decoder, *err*_*vt1*_ is the velocity error at time bin *t* using the single linear decoder (velocity decoder 1), *err*_*ft3*_ is the force prediction error at time bin *t* using the two-stage decoder, and *err*_*ft1*_ is the force error at time bin *t* using the single linear decoder (force decoder 1). *p*_*ve*_ and *p*_*fe*_ were then used to test if this two-stage decoder had an improved accuracy over the single linear decoder. Table 1 showed the flow charts for the three decoders.

**Table 1.**
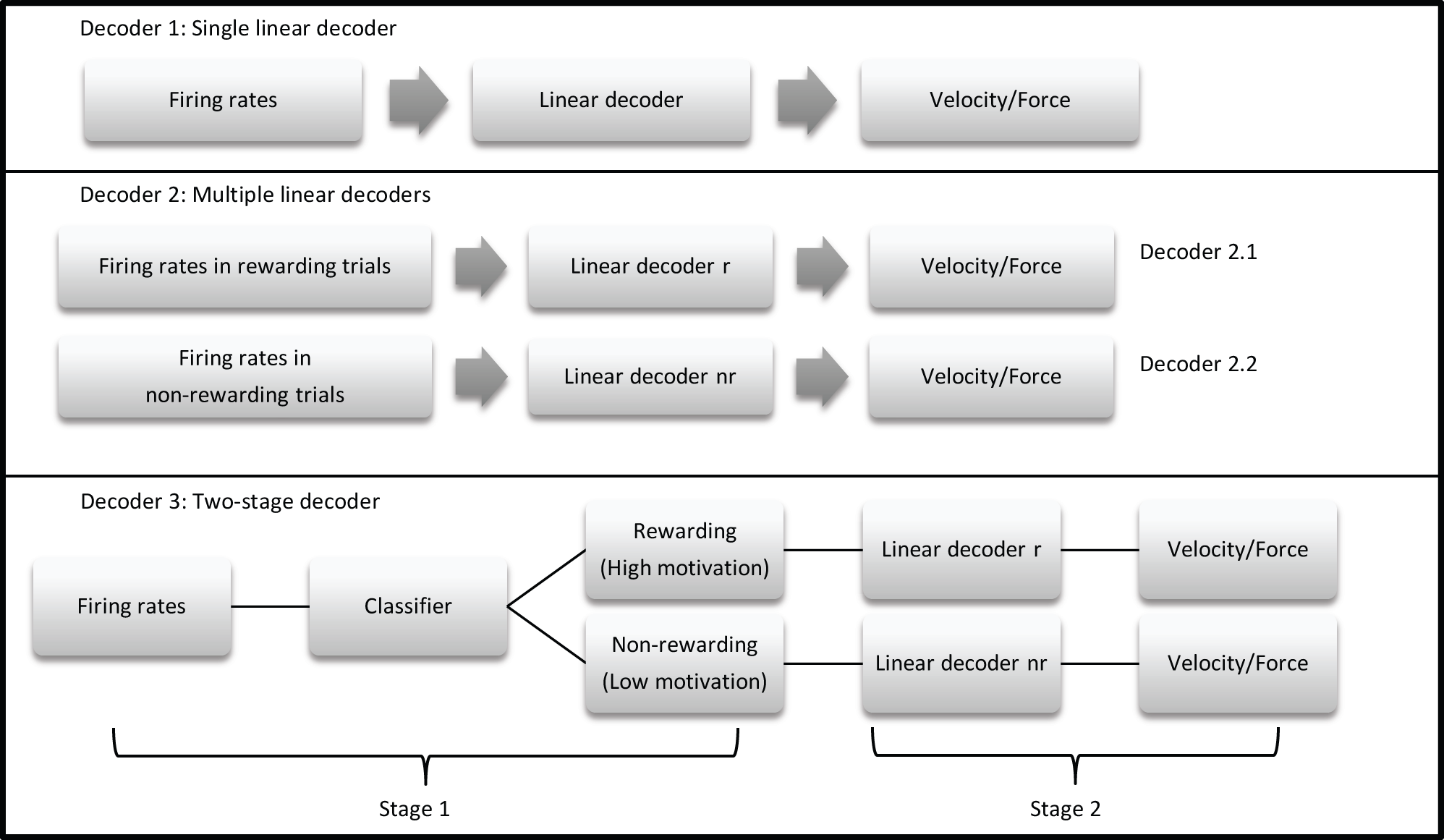
Three proposed decoders. Decoder 1 is a single linear decoder, decoding either velocity or force from M1 firing rates using a single linear decoder for all trials. Decoder 2 consists of two separate linear decoders, one for rewarding and one for non-rewarding trials. Decoder 3 is a two-stage decoder, which first classifies reward information from firing rate data then uses different decoders for different reward levels.

## 3 Results

The two NHP subjects in this work conducted two manual grip force sessions, and BMI sessions each. For the manual grip force task, NHP-P, 127 trials in session one with reward = R[0,1] and 156 trials in session 2 with R[0,3]. NHP-S 190 trials for session one in manual with R[0,1] and 161 trials in session two with R[0,3]. In addition, each NHP completed two BMI sessions. NHP-P 94 trials in session one with R[0,1] and 118 trials in session 2 with R[0,3]. NHP-S 82 trials for session one in BMI with R[0,1] and 177 trials in session two with R[0,3].

### 3.1 Directional tuning curves change based on reward level under BMI control

M1 units were used to decode movement information based on their tuning curves. The hypothesis was that for a given unit, tuning curve parameters would change based on the presence or absence of cued reward. For the BMI task (Fig. 2) a total of 87 units from monkey S and 102 units from monkey P were recorded. Of these, 52 units (60%) from monkey S and 42 units (41%) from monkey P had significant directional tuning (see Fig. 3 and 4). Of these significantly directionally tuned units we then asked if they showed significantly different tuning function parameters between the rewarding and non-rewarding trials. Note, that during the actual BMI task a single BMI decoder was used and the following analysis refers to offline analysis of those BMI experiments. We found that monkey S had 28 units (32%) and monkey P had 22 units (22%) with significantly reward-modulated preferred directions under BMI control. A larger percentage of units had significant reward modulation of their tuning curve model amplitude, with 48 units (55%) for monkey S and 39 units or (38%) for monkey P. These results are summarized in Fig. 4 with example units shown in Fig. 3.

**Figure 3.**
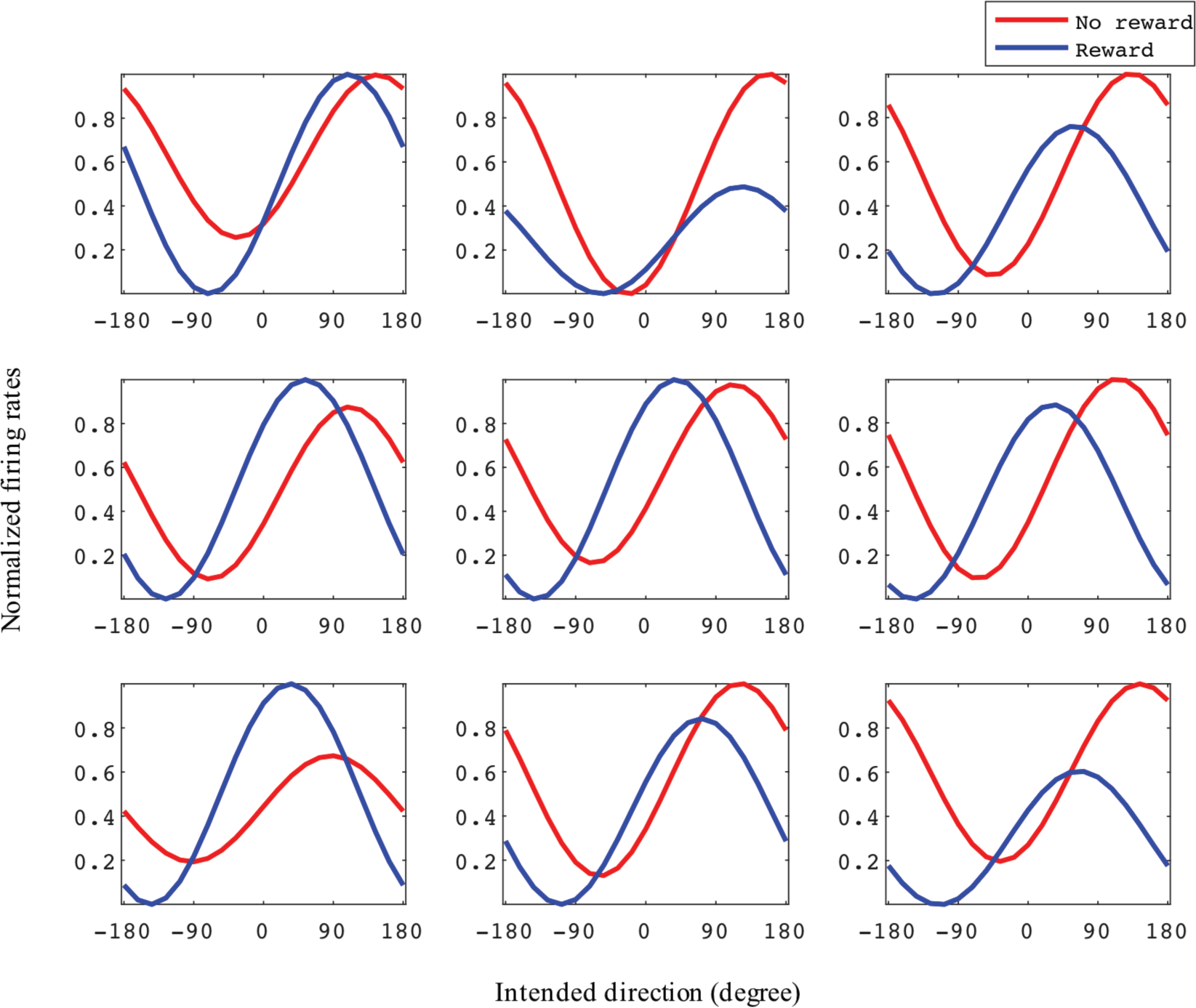
Example tuning curves for rewarding and non-rewarding trials for the BMI task. The x-axis is intended direction (degrees), and y-axis is normalized firing rates. Each subplot represents one unit. All example units were recorded from monkey S where the reward level for rewarding trials was 3.

**Figure 4.**
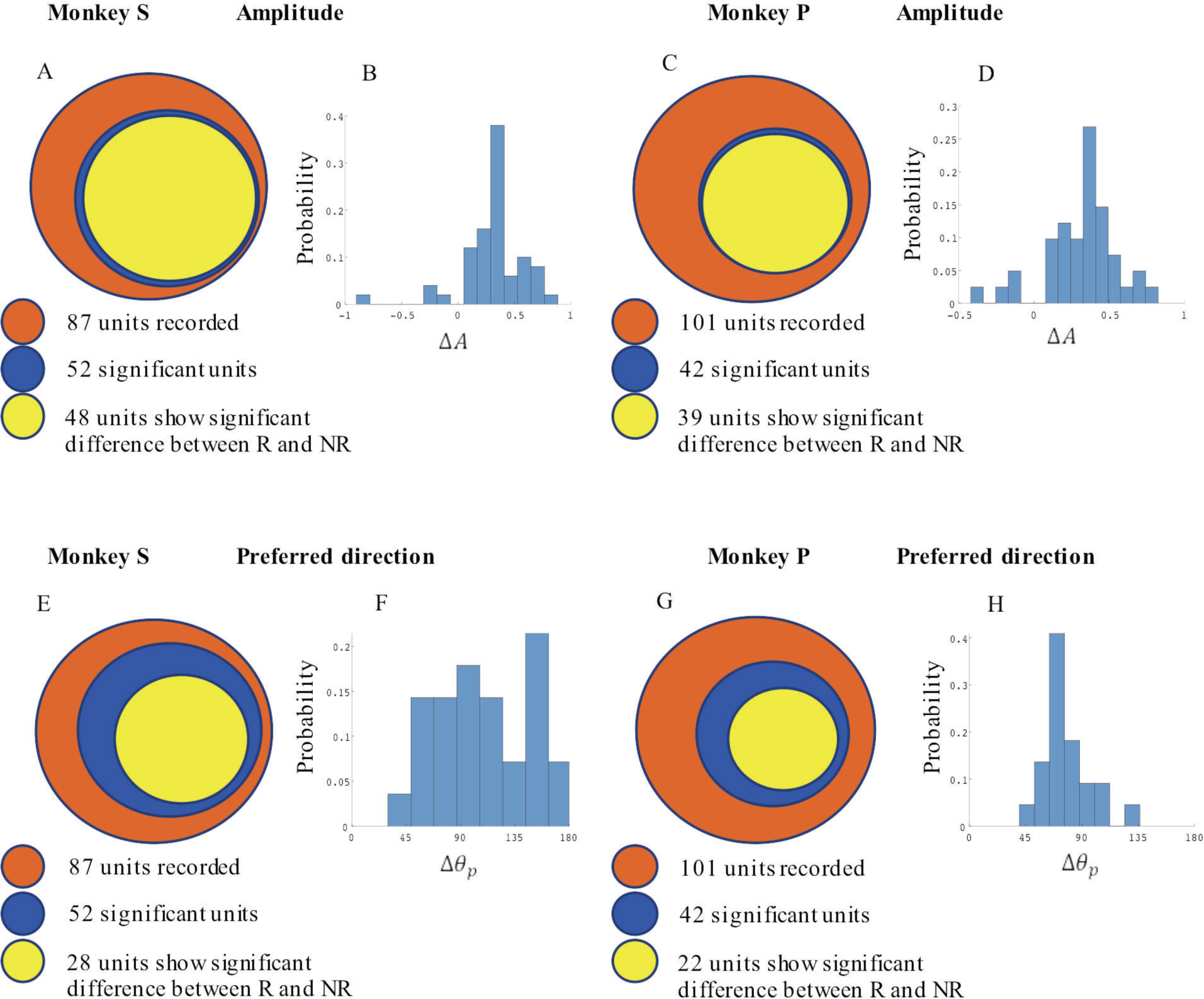
Statistical results for amplitude and preferred direction differences between rewarding (R) and non-rewarding (NR) trials in the BMI task. Monkey S **((A), (B), (E)** and **(F))** had a total of 52 significantly reward modulated units, and monkey P **((C), (D), (G)** and **(H))** had 42 units. **(A)** and **(C)** show the number of units which had significant changes in amplitude among all units and **(E)** and **(G)** show the preferred direction results. **(B)** and **(D)**show the distribution of ∆*A* for all units which had significant changes between R and NR. **(F)** and **(H)** show the distribution of ∆*θ_p_*. All units were recorded in blocks where reward levels were either zero or three.

In Fig. 3, we have plotted tuning curves for example units showing significant tuning functions that were also significantly reward modulated during the BMI task seen in Fig. 2. For units with a significant difference between rewarding and non-rewarding trials in either amplitude or preferred direction, the distribution of the changes in amplitude ∆*A* and change in preferred direction ∆*θ*_p_, are shown in Figure 4.1b and 4.1d for monkey S and Figure 4.2b and 4.2d for monkey P. The distribution of ∆*A* indicates that, on average, tuning curve amplitudes are larger for rewarding trials compared to non-rewarding trials.

### 3.2 Reward level modifies force tuning curves

In the manual grip force task, M1 units encode force and value. Figure 5 shows linear force tuning curves for six example units, fit using equations (4.1) and (4.2).

**Figure 5.**
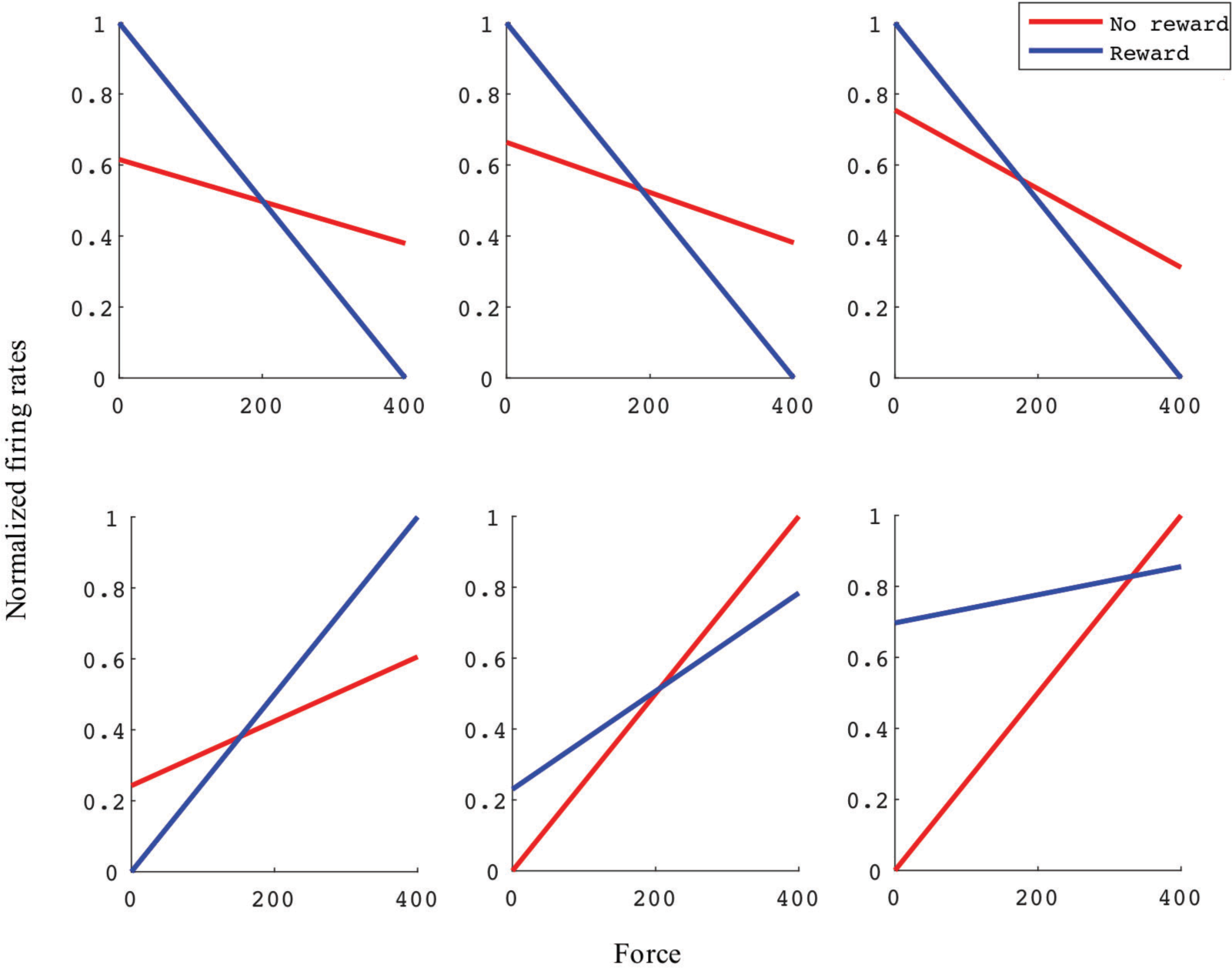
Linear force tuning curves between rewarding and non-rewarding trials for six example significant units. Both tuning curve characteristics, including slope and intercept, can change from rewarding to non-rewarding trials. X-axis is reading from force sensor. Y-axis is normalized firing rate. Each subplot represents one unit. All example units were recorded in monkey S M1 from an experimental block with reward levels of zero and three.

Significant force tuning was noted in 77 units (100%) in monkey S and 35 units (55%) in monkey P. Of these units with significant force turning, 31 units (40%) from monkey S and 10 units (16%) from monkey P were also significantly modulated by reward, having significantly different force turning curve slopes between rewarding and non-rewarding trials (see methods). Figure. 6 summarizes these results and the distribution of changes in force tuning slopes ∆*#x03B1;*, which are the normalized difference between rewarding and non-rewarding slopes (see Eq. 4.3).

**Figure 6.**
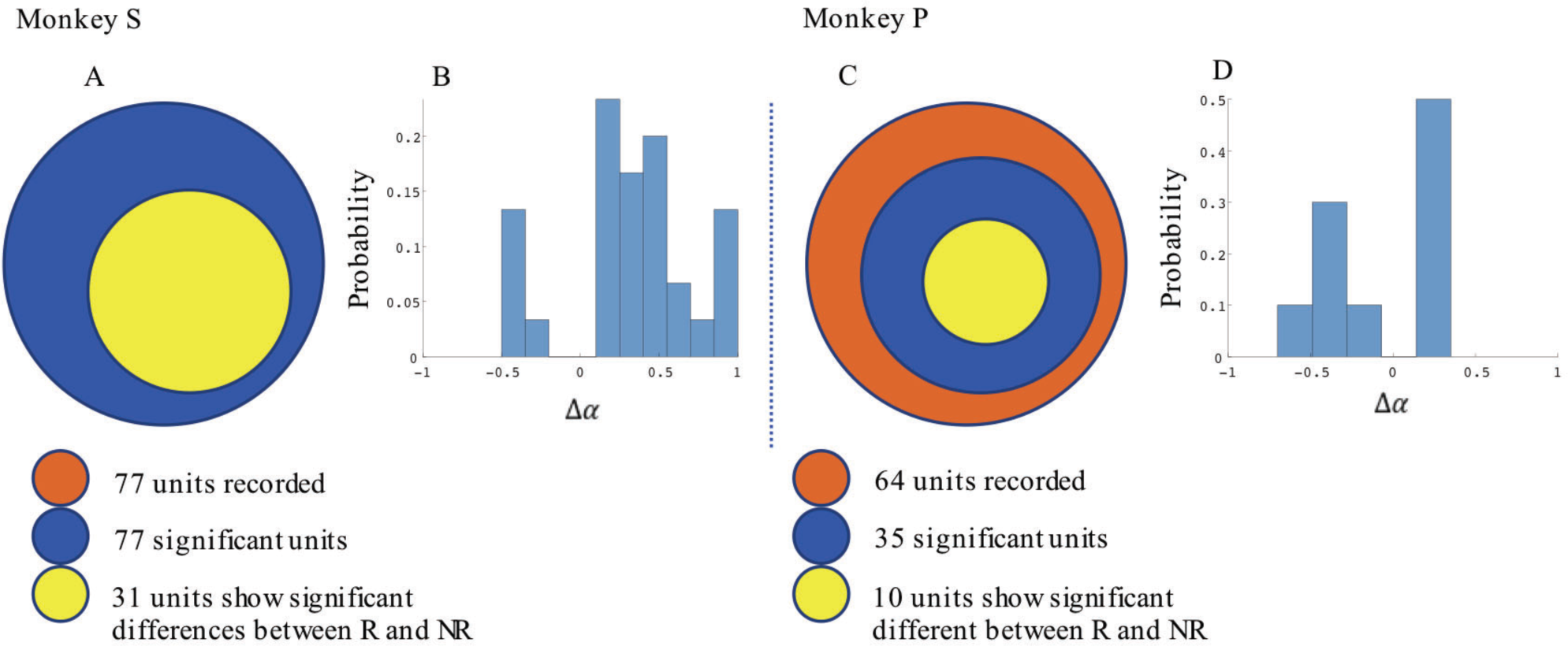
Statistical results for slope difference between rewarding and non-rewarding trials in the manual grip force task. **(A)** and **(B)** show results from monkey S and **(C)** and **(D)** are from monkey P. The number of units which have significant changes in their force tuning curve slopes and the distribution of ∆*α* for these units is shown in **(A)** and **(B)** for monkey S, and in **(C)** and **(D)** for monkey P.

### 3.3 Incorporating reward level improves force and movement decoding accuracy

The difference in directional tuning curves (Figs. 3 and 4) for individual units and the population suggest that units have different movement encoding for rewarding and non-rewarding trials, and that it may be helpful from a BMI decoding perspective to use models that allow for the influence of reward and motivation. Similarly, Figures 5 and 6 show that units change their force tuning curves based on reward level. This suggests that if two separate linear decoders are used for velocity or force decoding, one for rewarding and one for non-rewarding trials, decoding accuracy will be improved compared to using a single linear encoder. This dichotomy was used to build a decoder that treated rewarding and non-rewarding trials separately (see equations (5) and (6) in methods). Table 2 shows that decoding predictions were more accurate when we used the reward modulated decoders as compared to a single decoder (see methods). The percentage improvement in velocity decoding, *p*_*ve*_, and force decoding, *p*_*fe*_, was clear whether we used reward levles of zero and one, or zero and three (see methods). However, the percentage improvement was greater when the difference in trial value was greater, that is for the zero vs. three valued trials. The reward modulated velocity decoder produced a 22% - 28% error reduction compared to the single linear decoder (table 2, *p*_*ve*_). The reward modulated force decoder resulted in an error reduction of between 10% and 24% compared to the single force decoder (Table 2, *p*_*fe*_). These results demonstrate an improved decoding accruracy when rewarding and non-rewarding trials are treated separately, particularly when the value differences are greater. Also, from control groups results, the error reduction from decoder 2 are all greater than shuffled groups (*p*_*ve*_ or *p*_*fe*_ is larger than its corresponding 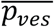 or 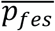), and that difference is significant (p<0.05, t-test). *p*_*ves*_ or *p*_*fes*_ distributions from control groups are normal (p<0.05, Jarque-Bera test). The results indicate that the decoding improvement did not because of decoder 2 have more parameters than decoder 1.

**Table 2.** Decoding accuracy was greater when multiple linear decoders corresponding to different reward levels were used (decoder 2) compared to the single linear decoder (decoder 1), for both velocity and force decoding. The control groups (shuffled decoder 2) do not show any improvement.

|  | Monkey S<br>(% error reduction) | Monkey P<br>(% error reduction) |
| --- | --- | --- |
| $R = 0$ or $1, p_{ve}$ | 23 | 22 |
| $R = 0$ or $1, \overline{p_{ves}}$ | 1.3 | 2.3 |
| $R = 0$ or $3, p_{ve}$ | 28 | 27 |
| $R = 0$ or $3, \overline{p_{ves}}$ | -1.1 | 0.29 |
| $R = 0$ or $1, p_{fe}$ | 11 | 10 |
| $R = 0$ or $1, \overline{p_{fes}}$ | -0.35 | 1.3 |
| $R = 0$ or $3, p_{fe}$ | 24 | 13 |
| $R = 0$ or $3, \overline{p_{fes}}$ | -1.9 | -2.2 |

### 3.4 Reward level is classifiable using post cue firing rates for cued BMI task

Results from Table 2, suggest that reward levels are useful information for BMI decoding. The classification accuracy and standard deviation across 10 Monte-Carlo repetitions for validation using post-cue firing rates to classify reward levels is shown in Table 3. For each Monte-Carlo repetition, 70% data were used for training and other 30% were used for testing. All classification accuracy is greater than chance (50%). These results demonstrate that reward level can be classified using firing rates. A subset of units using the best individual unit procedure (Leavitt et al 2017b) provides more accurate results than when all units were used.

**Table 3.** Reward level classification mean accuracy and standard deviation across 10 Monte-Carlo repetitions using post-cue firing rates and classifying between rewarding (R>0) and non-rewarding (R=0) trials.

| | Using all units<br>Mean accuracy ± s.d. | Using a subset of units<br>Mean accuracy ± s.d. |
| --- | --- | --- |
| $R = 0$ or $1$ | 68% ± 3.8% | 74% ± 1.6% |
| $R = 0$ or $3$ | 73% ± 2.7% | 78% ± 2.5% |

### 3.5 Two-stage decoder can improve decoding accuracy in offline test

Since the reward level can be classified using firing rates (table. 3), a two-stage decoder was made. Table 4 shows velocity decoding improvement *p*_*ve*_ and force decoding improvement *p*_*fe*_ for the offline test of the two-stage decoder using 5-fold cross-validation for both monkeys. Here the reward level is decoded during the first stage from the neural data and is used to pick the equations for the second stage. It should be noted that the percent improvements *p*_*ve*_ and *p*_*fe*_ are greater than 0 for all cases, therefore the two-stage decoder is more accurate than the single linear decoder.

**Table 4.** Two-stage decoder improvement over decoder 1. For both velocity and force decoding, two-stage decoder reduced decoding error.

|  | Monkey S<br>(% error reduction) | Monkey P<br>(% error reduction) |
| --- | --- | --- |
| $R = 0$ or $1, p_{ve}$ | 14 | 15 |
| $R = 0 \text{ or } 3, p_{ve}$ | 8 | 7 |
| $R = 0 \text{ or } 1, p_{fe}$ | 7 | 7 |
| $R = 0 \text{ or } 3, p_{fe}$ | 10 | 7 |

## 4 Discussion

In the current work, NHP subjects controlled either grip force manually or reaching kinematics with a BMI. In both cases the subjects controlled a simulation of an anthropomorphic robotic arm or hand to reach, grasp and transport target objects. Each trial was cued as to the reward value the subject would receive for making the correct movement. In this work, we found that M1 unit activity was modulated by cued reward level in both the manual grip force task and the BMI kinematics control task. In both of these tasks the neural tuning functions, force tuning and kinematic tuning, were significantly modulated by the level of expected reward in blocks of trials where cued reward was (zero or one), or (zero or three). Our results indicate that reward influences motor related encoding in both manual and BMI tasks. When we explicitly took the influence of motivation/value into consideration, our linear decoding models predicted with significantly higher accuracy. Having a more predicative linear decoding model is important because the most successful BMI systems use such linear models at their core, such as within a Kalman framework, or simply use the output of a linear model that decodes neural rate into movement parameters. A more accurate linear decoding model should lead to a more controllable BMI. Regardless of the BMI control aspects of this work, the basic neuroscience stands, which is that expected reward modulated M1 motor related tuning functions. We show in the current work and in our previous work (Marsh et al 2015, McNiel et al 2016a, McNiel et al 2016b) that the cued reward level in a task is classifiable using post-cue firing rates, therefore, it should be possible to build a classifier to determine the motivation/reward level before sending the neural activity through the appropriate BMI decoder. We have recently found that M1 activity in NHPs is also predictive of un-cued reward levels if these levels are predictable (Tarigoppula et al 2017). This indicates that the system we are proposing, which uses a neural critic/classifier to switch between BMI decoders could have relevance past the laboratory setting with learned cues.

It can be seen from Table 3 that classification accuracy is higher when the difference between the reward levels is larger, that is between trials with cued reward of (0 vs. 3) as compared to trials with cued reward of (0 vs. 1). This result indicates that neural firing rates are more separable when there is a higher cued value difference. The same conclusion can also be inferred from Table 2, where both percent improvement for velocity decoding (*p*_*ve*_) and for grip force decoding (*p*_*fe*_) from trials with reward levels (0 vs. 3) are larger than (0 vs. 1). We have recently found that at least some M1 units code reward level in a linear manner in agreement with the above results (Tarigoppula et al 2017). Table 2 shows decoding results for separate linear decoders where the reward level was specified by the experimenter as an additional input to the decoder. Table 4, on the other hand, shows decoding results of the two-stage decoder, where the reward level is automatically classified in the first stage using a kNN algorithm. The only difference between these two versions is whether the experimenter has to provide added information, or if the system can do it autonomously. Since the reward level classifier for the two-stage decoder was not 100% accurate, the *p*_*ve*_ and *p*_*fe*_ from Table 4 are lower than the values in Table 2. It can be seen that automatic classification of reward level comes at the cost of decreased decoding accuracy as expected, indicating the value of a highly accurate classifier if this strategy is to be meaningful. We have previously shown that one can also obtain cued reward level information from local field potentials (Marsh et al 2015, Tarigoppula et al 2017) that are concurrently obtained when recording single units. Recently we have obtained up to 98% classification accuracy by combining features that include both spike related information and LFP based information (An et al 2017), and we will incorporate such information in our online neural critic in future work.

From the current work, it is clear that neural firing rates in M1 do not only contain pure movement related information. As shown in this and in previous work, we have made use of reward information in a controlled environment by incorporating reward levels to develop a more robust and accurate decoder. However, in a more realistic scenario the number of reward levels may not be known, and there may be other unknown factors encoded in M1. It will be more meaningful to have a dynamic decoder that is able to filter out the reward based information for any level of reward and reject the effect of other such “non-movement” variables to obtain more purely movement relevant information for BMI control. This way we can build a more stable BMI decoder able to work in a more complex, naturalistic environment.

## 3 Author Contributions

YZ, JPH and JTF conceived the research. YZ and JPH conducted the experiments. YZ conducted the analysis. YZ, JPH, JK and JTF wrote the paper.

## Acknowledgments

We would like to thank SUNY Downstate DMC for all their help with the NHPs.

